# «*In vitro* and *in vivo* combination of lytic phages and octapeptin OPX10053 against β-lactamase-producing clinical isolates of *Klebsiella pneumoniae*»

**DOI:** 10.1101/2023.03.15.532768

**Authors:** Olga Pacios, Lucia Blasco, Ines Bleriot, Laura Fernández-García, María López, Concha Ortiz-Cartagena, Antonio Barrio-Pujante, Felipe Fernández Cuenca, Belen Aracil, Jesús Oteo-Iglesias, Karl A. Hansford, María Tomás

**Affiliations:** Microbiology Department-Research Institute Biomedical A Coruña (INIBIC); Hospital A Coruña (CHUAC); University of A Coruña (UDC), A Coruña, Spain; Study Group on Mechanisms of Action and Resistance to Antimicrobials (GEMARA) on behalf of the Spanish Society of Infectious Diseases and Clinical Microbiology (SEIMC), Madrid, Spain; Clinical Unit of Infectious Diseases and Microbiology, Hospital Universitario Virgen Macarena, Institute of Biomedicine of Seville (University Hospital Virgen Macarena/CSIC/University of Seville), Seville, Spain; Reference and Research Laboratory for Antibiotic Resistance and Health Care Infections, National Centre for Microbiology, Institute of Health Carlos III, Majadahonda, Madrid, Spain; Centre for Superbug Solutions, Institute for Molecular Bioscience, University of Queensland, St Lucia, QLD. 4072. Australia; CIBER de Enfermedades Infecciosas (CIBERINFEC), Instituto de Salud Carlos III, Madrid, Spain

**Keywords:** Phages, octapeptin, synergy, *Klebsiella pneumoniae*

## Abstract

**Background:** novel approaches to treat *Klebsiella pneumoniae* infections are desperately needed, such as the use of rationally designed combination therapies.

**Objectives:** to evaluate the *in vitro* and *in vivo* therapeutic potential of lytic phages against *K. pneumoniae* in combination with octapeptin, a promising class of lipopeptides with broad spectrum Gram-negative activity.

**Methods:** we determined the MICs to twenty-two lipopeptide compounds and chose one octapeptin (OPX10053) for evaluation of potential synergism in combination with lytic phages using checkerboard assays, optical density growth curves and time-kill (CFU enumeration). Toxicity and efficacy *in vivo* assays were conducted on *Galleria mellonella* larvae.

**Results:** this study reports the synergy found *in vitro* between the octapeptin OPX10053 and two lytic phages previously characterized by our research group (vB_KpnM-VAC13 and vB_KpnM-VAC66) against clinical isolates of *K. pneumoniae*. This synergy was validated by the FIC index, OD growth curves and time-kill assay when OPX10053 was added following 4 hours of phage exposure. Preliminary evaluation of toxicity revealed that OPX10053, even at subinhibitory concentrations and in phage combinations, exerts a toxic effect on larvae, which requires further investigation.

**Conclusions:** The *in vitro* application of lytic phages in combination with octapeptin OPX10053 showed synergistic activity. Exposure of *G. mellonella* to the lytic phages was well tolerated, whereas combination treatment with subinhibitory concentrations of OPX10053 did not attenuate toxicity. Even so, this innovative approach of combining lytic phages could open the door to some interesting associations between chemically synthesized drugs and biological entities. Sequential or simultaneous application alongside time, dosing and stewardship warrants further research.

## Introduction

*Klebsiella pneumoniae* is an increasingly worrisome opportunistic pathogen that causes severe to life-threatening infections in the urinary tract, lungs, blood and soft tissues of immunocompromised patients ^1, 2^. *K. pneumoniae* infections are alarming because numerous strains are resistant to many contemporary antibiotics, creating a scenario reminiscent of the pre-antibiotic era ^3^. In particular, carbapenem-resistant *K. pneumoniae* represents a serious concern and is responsible for 600 deaths and 9000 infections in the United States annually ^4^. Persistent strains of this species display a plethora of altered mechanisms to overcome antibiotic treatment by entering a dormant state, leading to chronic and recurrent infections that are extremely difficult to eradicate ^5^. Furthermore, clinical *K. pneumoniae* strains easily acquire antibiotic resistance plasmids, form biofilms and overexpress efflux pumps ^6, 7^.

To combat this looming health threat, novel approaches to antimicrobial treatment are desperately needed. Antimicrobial peptides (AMPs) are noteworthy as they have shown low rates of resistance and display activity against many multidrug-resistant (MDR) isolates. One of the best-studied peptide antibiotics are polymyxins, which have been reserved as a last-resort treatment, albeit suffering from nephrotoxicity and neurotoxicity ^8-10^. Efforts have been made to generate lipopeptide analogues with improved toxicity profiles ^11, 12^.

One attractive type of lipopeptide class are the octapeptins, which differ from polymyxins by possessing two fewer amino acids in the exocyclic tail with inverted stereochemistry in the exocyclic diaminobutyric acid (Dab) residue ^13^. Both classes retain a heptapeptide core, but in the case of octapeptins this is bound to a lipophilic acyl monopeptide tail (β-hydroxy fatty acid), exemplified by octapeptin C4, one of 18 congeners within this natural product class (Figure 1) ^14^. Despite the structural similarities of both classes, octapeptins are of considerable interest due to their intrinsic *in vitro* activity against polymyxin-resistant Gram-negative bacteria ^15, 16^. Moreover, the recent demonstration that repeated exposure of a clinical isolate of *K. pneumoniae* to sub-lethal doses of polymyxin or octapeptin C4 over 20 days of daily passage leads to divergent levels of resistance development (1000-fold increase in MIC for polymyxin vs 5-fold increase for octapeptin) with no cross-resistance supports the further development of the octapeptins as a unique class of antibiotics ^17^.

**Figure 1:**
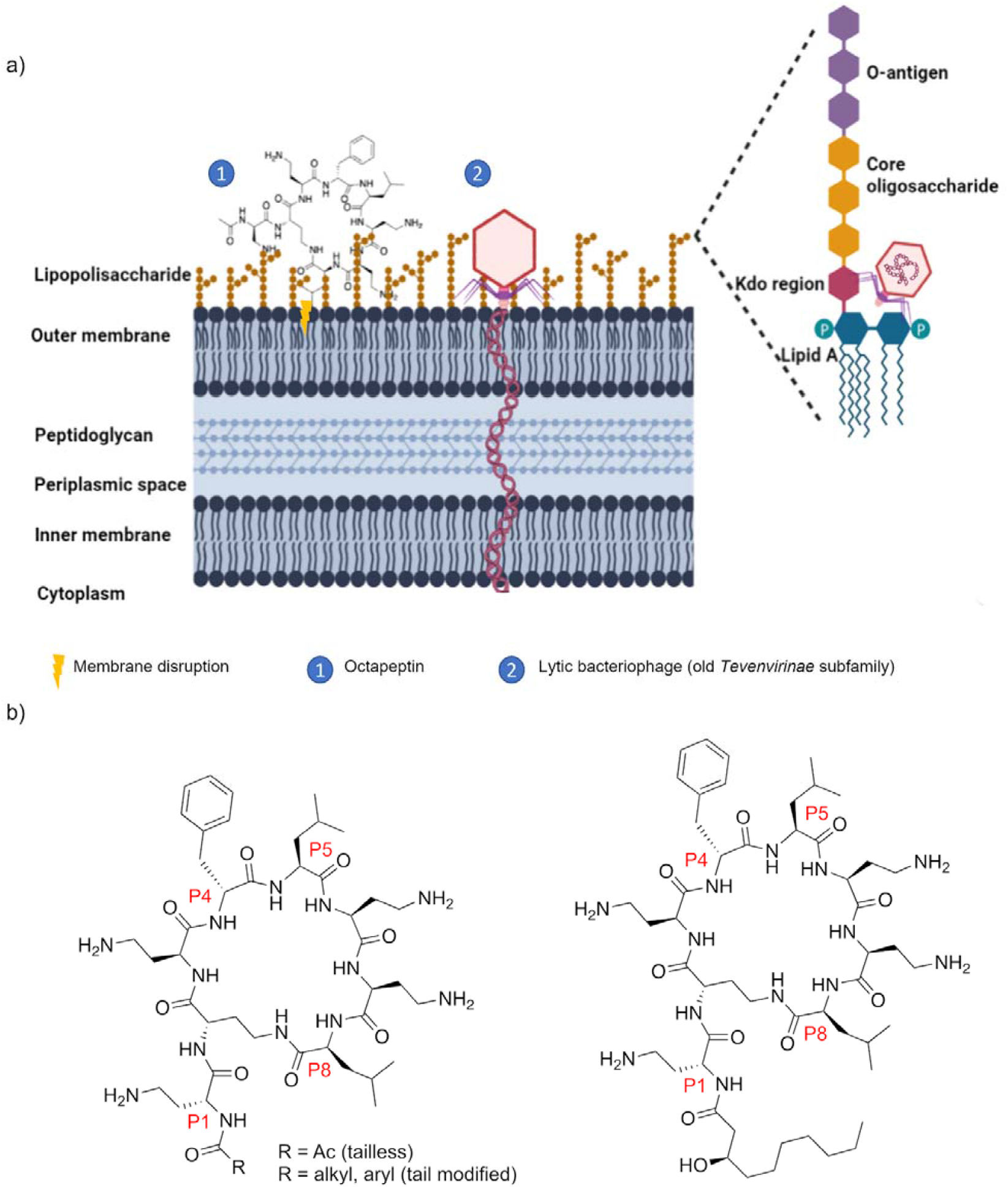
a) Proposed mechanism of action for synergy between this drug-biologic combination. b) Chemical structures of polymyxins and octapeptins.

Lipopeptide antibiotics such as polymyxin possess complex structure-activity and structure-toxicity interrelationships, with nephrotoxicity and acute toxicity major contributors to pipeline attrition. Strategies to ameliorate the toxic liabilities of polymyxin, including medicinal chemistry optimisation and the application of truncated versions of polymyxins that lack intrinsic antibacterial activity but facilitate entry of co-dosed partner antibiotics by permeabilising the outer bacterial membrane, have identified promising pre-clinical candidates ^11, 12, 18^. Currently, little is known about the toxicity of octapeptins, although preliminary reports suggest that, in mice, octapeptin C4 and octapeptin B5 exhibit reduced nephrotoxicity and/or acute toxicity compared to polymyxin ^16, 19^. In this context, we considered a strategy in which the co-dosing of octapeptin in combination with lytic phages might provide a cooperative killing effect against hard-to-treat *K. pneumoniae* isolates, characterised by the use of sub-inhibitory concentrations of octapepin as a means to attenuate potential toxicity ^20-23^ (Figure 1).

Phages (bacteriophages) are self-replicating, biological entities that specifically infect their bacterial hosts, minimising the dysbiosis of the normal microbiota ^24^. Phages displaying a lytic cycle are the ones prioritised for therapy, though interesting studies have shown the potential of re-engineering lysogenic phages to produce novel lytic phages ^25^.

Our group recently characterized and compared two lytic phages (vB_KpnM-VAC13 and vB_KpnM-VAC66) with a broad lytic spectrum against *K. pneumoniae* clinical isolates ^26^. These phages belong to the “old” *Tevenvirinae* subfamily, proposed to target the deep sugar motifs in the LPS core as secondary receptors ^27^. With this in mind, we aimed to search for synergistic interactions between chemically synthesized octapeptins and the lytic phages vB_KpnM-VAC13 and vB_KpnM-VAC66 in four heterogeneous clinical isolates of *K. pneumoniae* and one reference strain (Table 1). Herein, we report the minimal inhibitory concentrations (MIC) of 21 octapeptin analogues against five strains of *K. pneumoniae* (K3320, K3324, K3325, K2534 and ATCC10031). From this pool, we identified the octapeptin analogue OPX10053, which was further characterised *in vitro* and *in vivo*. Reduction in viability was determined and compared to the monotherapies and the non-treated control, and efficacy was assessed in *Galleria mellonella* larvae infection model.

**Table 1:** Genomic and epidemiologic characteristics of *K. pneumoniae* isolates used in this study. ST: sequence type; OXA-245: oxacillinase carbapenemase; KL: K-locus

| *K. pneumoniae* strain | ST | β-lactamase genes (*bla*) | Hospital | Biological origin | Capsular type |
| --- | --- | --- | --- | --- | --- |
| **ATCC 10031** |  | - | Commercial | Commercial | No genome publicly available |
| **K3320** | 163 | SHV-36 | Virgen Macarena (Spain) | Blood | KL139 |
| **K3324** | 542 | SHV-1 | Virgen Macarena (Spain) | Blood | KL8 |
| **K3325** | 42 | SHV-1 | Virgen Macarena (Spain) | Blood | KL64 |
| **K2534** | 437 | OXA-245, CMY2, SHV-182 | National Centre of Microbiology (Spain) | Rectal | KL36 |

## Material and methods

### Bacterial strains, lytic phages and growth conditions

Clinical strains of *K. pneumoniae* K3320, K3324 and K3325 were isolated in the University Hospital Virgen Macarena (Sevilla, Spain), whereas K2534 was isolated in the National Centre of Microbiology (Madrid, Spain). The lytic phages vB_KpnM-VAC13 and vB_KpnM-VAC66 were isolated from sewage water in Valencia (Spain) and were further characterized and studied in previous works by our group ^26^. Luria-Bertani (LB) broth was used to grow the overnight cultures and LB supplemented with 1mM of CaCl_2_ was used when cultures were infected with phages, to enhance adsorption ^28^. For visualization of lysis plaques, TA (1% tryptone, 0.5% NaCl, 1.5% agar) and semi-solid Soft (1% tryptone, 0.5% NaCl, 0.4% agar) media supplemented with 1mM CaCl_2_ were used.

### Chemically synthesized octapeptins used in this study

A series of 21 lipopeptides, comprising 4 polymyxin and 17 octapeptin analogues, were obtained from the University of Queensland. The polymyxin compounds included polymyxin B, SPR-206 ^12^, FADDI-287 ^29^, and QPX-9003 ^11^. Within the octapeptin series, 7 possessed a 3-hydroxydecanoic acid tail, (designated “octapeptins”), 6 possessed different fatty acyl tail substituents, (designated “tail modified”), and the remaining 4 were capped at the N-terminus with an acyl group (designated “tailless”), combined with various ring modifications at ring positions P8, P5, P4 and P1 (Figure 1b).

### MIC determinations to peptide compounds and lytic phages

The MIC of every peptide compound used in this study was assessed by the broth microdilution method following the EUCAST guidelines ^30^. Briefly, 2-fold serial dilutions of the peptides were tested in concentrations ranging from 32 to 0.125μg/mL, and with a starting inoculum of 5·10^5^CFU/mL. 384-well flat-bottom plates and sterile Cation-Adjusted Muller Hinton Broth were used, with a final volume of 50μL.

To assess the inundation threshold of the lytic phages (the phage titre required to cause a decrease in the bacterial population), 10-fold dilutions were performed in SM buffer (20mM Tris-HCl pH=7.5, 1mM MgSO_4_ and 10mM NaCl) in 96-well polystyrene plates, based on other works ^31, 32^. As previously mentioned, we used LB+1mM CaCl_2_. Every bacterial strain was inoculated at a final concentration of 5·10^5^ CFU/mL per well. The OD_600 nm_ was measured every hour to assess at which time-point phage resistance would arise.

In both cases, plates were statically incubated at 37°C in the plate reader instrument Infinite® M1000 i-Tecan™ and the MIC was defined as the minimal concentration of each compound or lytic bacteriophage in which no growth was visible after 18h of incubation. Every MIC determination was performed in technical duplicates and repeated in three independent experiments.

### Checkerboard assay

Among all the peptide compounds, OPX10053 was chosen for further experiments. This peptide was 2-fold serially diluted in 50μL of LB+1mM CaCl_2_, ranging from 32 to 0.03 μg/mL along the X-axis of 96-well microtiter plates. 25μL of either vB_KpnM-VAC13 or vB_KpnM_VAC66 were added to the columns at concentrations ranging from 10^9^ to 10^5^ plaque-forming units (PFU)/mL, 10-fold diluted in SM buffer. Overnight cultures of *K. pneumoniae* were diluted 1:100 in LB broth and incubated until OD_600 nm_ =0.5-0.8, then 25μL were added to the wells at a starting inoculum of 5×10^5^CFU/mL. The fractional inhibitory concentration (FIC) for octapeptin OPX10053 was calculated using the following formula: FIC_OPX10053-Phage_ = MIC_OPX10053_ in presence of phage / MIC_OPX10053_ alone, as described in other works ^31^. The plates were statically incubated at 37°C then the OD_600 nm_ was checked at 18h using the Infinite® M1000 i-Tecan™ plate reader.

### Resazurin viability assay

Checkerboard plates were incubated with 0.002% of resazurin, a cell-permeable redox indicator that is reduced to a pink resorufin product within viable, actively metabolic cells. 10μL of water-diluted resazurin were added to each well. Plates were incubated for 2h at 37°C until colouration was visible. Blue-coloured wells indicate an absence of metabolically active cells, whereas pink wells were indicators of bacterial metabolism.

### Optical density growth curves in presence of lytic phages and the peptide compound sequentially applied

Overnight cultures of *K. pneumoniae* K3324, K2534 and ATCC 10031™ were 1:100 diluted in LB broth and incubated at 37°C for 2h until the culture reached an early exponential phase. This assay was performed in 96-well microtiter plates, using LB+1mM CaCl_2_. Phages were inoculated at 10^9^PFU/mL and, for the first 4h of incubation, cells were exclusively exposed to them. After that, the octapeptin OPX10053 was added at 2μg/mL final concentration for the clinical strains K2534 and K3324 and at 1μg/mL for the reference strain ATCC 10031 (1/2 MIC, respectively), to every well except to the growth controls and the only-bacteriophage control group. The bacterial inoculum was 5·10^5^CFU/mL per well.

### Time-kill assay in presence of lytic phages and the peptide compound sequentially applied

A spotting assay was performed to elucidate the dilutions in which individual colonies were accountable. At the desired time-points (0, 4, 6 and 24 hours post-infection -hpi-), 50μL of each well were transferred from the time-kill plate to the first row of the dilution plate, containing 50μL of charcoal suspension to inactivate the compound (25mg/mL) in row A, and 90μL of 0.9% sterile saline in rows B-H. After the initial 1:2 dilution, 10μL were serially transferred with a multichannel pipette, then 10μL of every dilution and condition was spotted onto large Petri dishes containing LB-agar. The following day, we homogeneously streaked 10μL in conventional-size LB-agar plates and calculated the CFU/mL considering dilution factors relating to the charcoal inactivation, the plated dilution and the 10μL volume streaked onto each plate.

### Toxicity and efficacy of combinations between lytic phages and the octapeptin OPX10053 in the *G. mellonella in vivo* model

*G. mellonella* larvae were acquired from DnatEcosistemas®, Spain. Only healthy larvae lacking dark spots were chosen, randomly allocated into groups (n=15) and kept in dark and starvation conditions in Petri dishes at 15°C for at least 24h prior to use. 10μL of the inoculum, phage and/or peptide solutions were injected in the last left proleg using a Hamilton micro-syringe. The inoculum was incubated overnight at 37°C at 180rpm, centrifuged (4000g, 15min) and washed 3 times with saline. Suspensions containing 10^8^CFU/mL (K3324), 10^9^CFU/mL (K2534), the phages or the OPX10053 for the toxicity assay were inoculated and, 90min after infection, the treatment groups were injected via the last right proleg with either 10μL of the lytic phages and/or the OPX10053 at ½ MIC (2μg/mL). The phages were previously purified using an Amicon® Centrifugal Filter Unit, following the protocol described in ^33^, and inoculated at MOI=1. We were unable to maintain the sequential approach performed *in vitro* (in which bacteria were exposed only to phage for 4h and the peptide was added later), since the larvae injected with sterile saline buffer 3 times showed high mortality (data not shown).

## Results

### MIC determination to peptide compounds

The MIC of twenty-two peptide compounds was assessed by the broth microdilution method and reported in Table 2. Meropenem and polymyxins (Pmx) were included as controls, and the peptide compounds were categorized into classes according to their chemical structure: the tailless compounds are devoid of the fatty acid tail, the octapeptins maintain a highly similar structure to the natural products, and the tail-modified compounds contain additional diversity within the fatty acyl tail (Figure 1b). All the strains were susceptible to the polymyxins (MIC=0.25μg/mL) and to meropenem, except for the clinical isolate K2534, which harbours a carbapenemase OXA-245 that confers resistance to carbapenems (MIC>2μg/mL). K3325 is an imipenem-persister strain, which leads to the assumption that other carbapenems such as meropenem would not be suitable for its treatment ^34^. Among all the peptide compounds, the tailless were inactive at the highest concentration tested (MIC values≥16 μg/mL), whereas the remainder of the lipopeptides exhibited variable MIC values ranging from 1 to 32μg/mL. Of note, octapeptin OPX10053 consistently showed relatively low MIC values for all the isolates (2-4 μg/mL), warranting further investigation.

**Table 2:** MIC values of the peptide compounds tested over the five *K. pneumoniae* strains, grouped according to their chemical properties.

|  | Carbapenem | Pmx |  |  |  | Tailless |  |  |  | Octapeptins |  |  |  |  |  |  | Exotic |  |  |  |  |  |
| --- | --- | --- | --- | --- | --- | --- | --- | --- | --- | --- | --- | --- | --- | --- | --- | --- | --- | --- | --- | --- | --- | --- |
|  | Meropenem | 636 | 9490 | 10041 | 9476 | 8980 | 9833 | 10030 | 9222 | 6442 | 631 | 5561 | 9295 | 9292 | 9628 | 9630 | 10050 | 10049 | 10057 | 9872 | 10053 | 9908 |
| ATCC | 0.23 | 0.25 | 0.25 | 0.25 | 0.25 | 16 | >32 | >32 | 16 | 2 | 4 | 2 | 1 | 1 | 1 | 1 | 1 | 4 | 2 | 1 | 2 | 2 |
| K3320 | 0.23 | 0.25 | 0.25 | 0.25 | 0.25 | >32 | >32 | >32 | 32 | 8 | 8 | 8 | 8 | 8 | 8 | 8 | 8 | 16 | 8 | 8 | 4 | 8 |
| K3324 | 0.23 | 0.25 | 0.25 | 0.25 | 0.25 | >32 | >32 | >32 | >32 | 16 | 16 | 16 | 16 | 16 | 8 | 8 | 32 | 32 | 32 | 16 | 4 | 32 |
| K3325 | 0.23 | 0.25 | 0.25 | 0.25 | 0.25 | 32 | >32 | >32 | >32 | 8 | 8 | 8 | 8 | 8 | 8 | 8 | 16 | 16 | 16 | 16 | 4 | 16 |
| K2534 | 7.25 | 0.25 | 0.25 | 0.25 | 0.25 | >32 | >32 | >32 | >32 | 16 | 16 | 16 | 16 | 8 | 16 | 8 | 8 | 32 | 32 | 8 | 4 | 16 |

MIC values (µg/mL) of the peptide compounds tested on *K. pneumoniae* strains. Pmx: polymyxins; tailless compounds: polymyxins derivatives without fatty acid tail; octapeptins: polymyxins derivatives showing slight amino acid changes; exotic compounds: polymyxins derivatives with other substitutions.

### Inundation threshold: “MIC” determination to lytic phages

To comprehensively assess the infection kinetics for each *K. pneumoniae* isolate by both phages, the OD_600 nm_ was measured every hour to have a better understanding of the precise time-point at which the bacterial resistance starts. We observed that phages were effective at cell lysis at the highest titre with no development of resistance until the titre was reduced from 4 hpi onwards, after which the OD_600_ increased dramatically (Figure 2).

**Figure 2:**
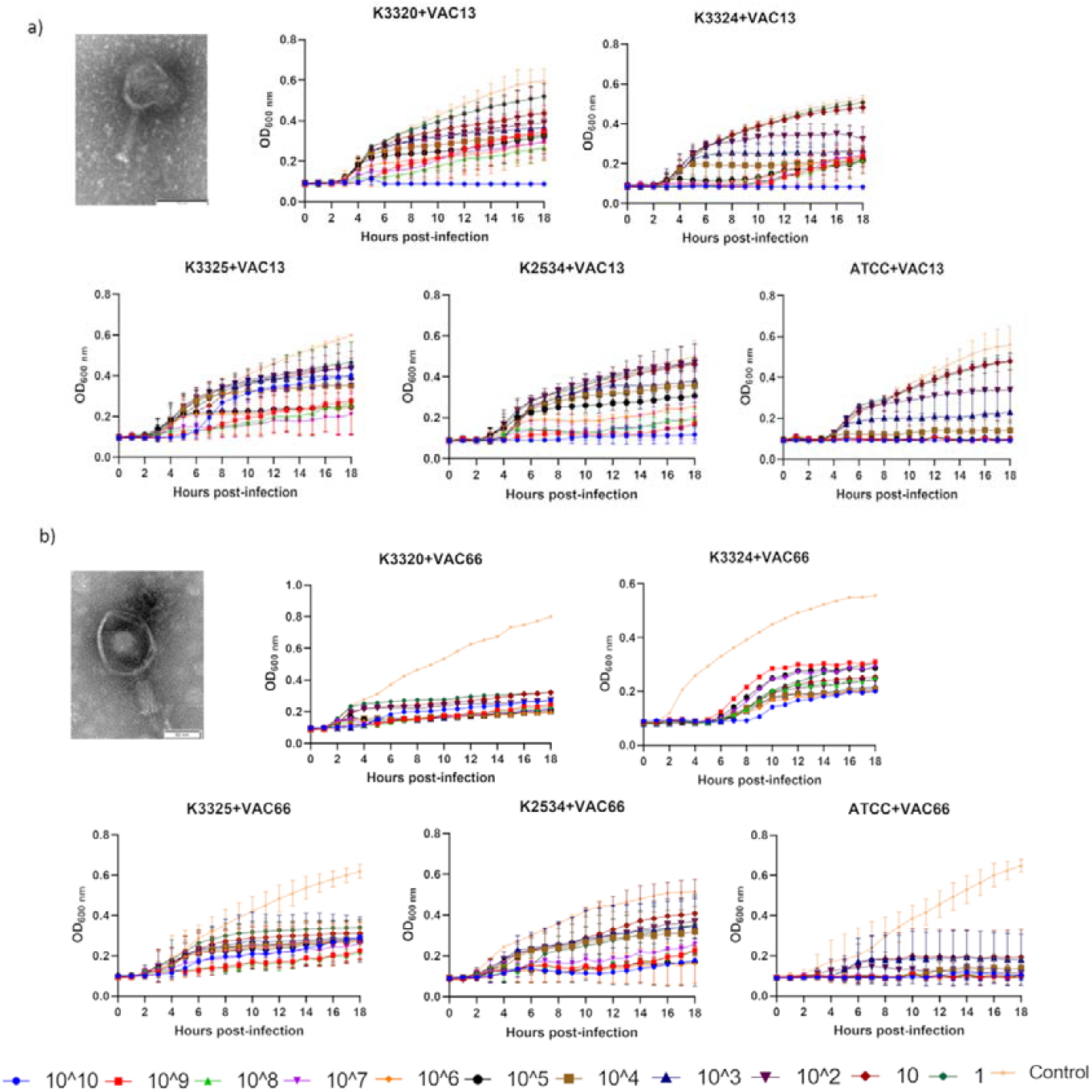
Kinetics of infection of *K. pneumoniae* isolates by the lytic phage vB_KpnM-VAC13 (a) and vB_KpnM-VAC66 (b) during time. Numbers represent the concentration of each phage in PFU/mL, and control means absence of phage.

### Checkerboard assays: FIC calculation

Following the standardized guidelines ^35^, a ≥2-fold reduction in the MIC to the combination compared to the MIC_OPX10053_ alone was considered a significant change in the bacterial strain susceptibility to the compound (FIC≤0.5). K3324, K2534 and ATCC 10031 exhibited an increase in susceptibility to the peptide OPX10053 in presence of phages, so they were selected for further assays. Resazurin staining of a representative checkerboard microtiter plate, and the plate layout together with the values obtained for each strain are reported in Figure 3.

**Figure 3:**
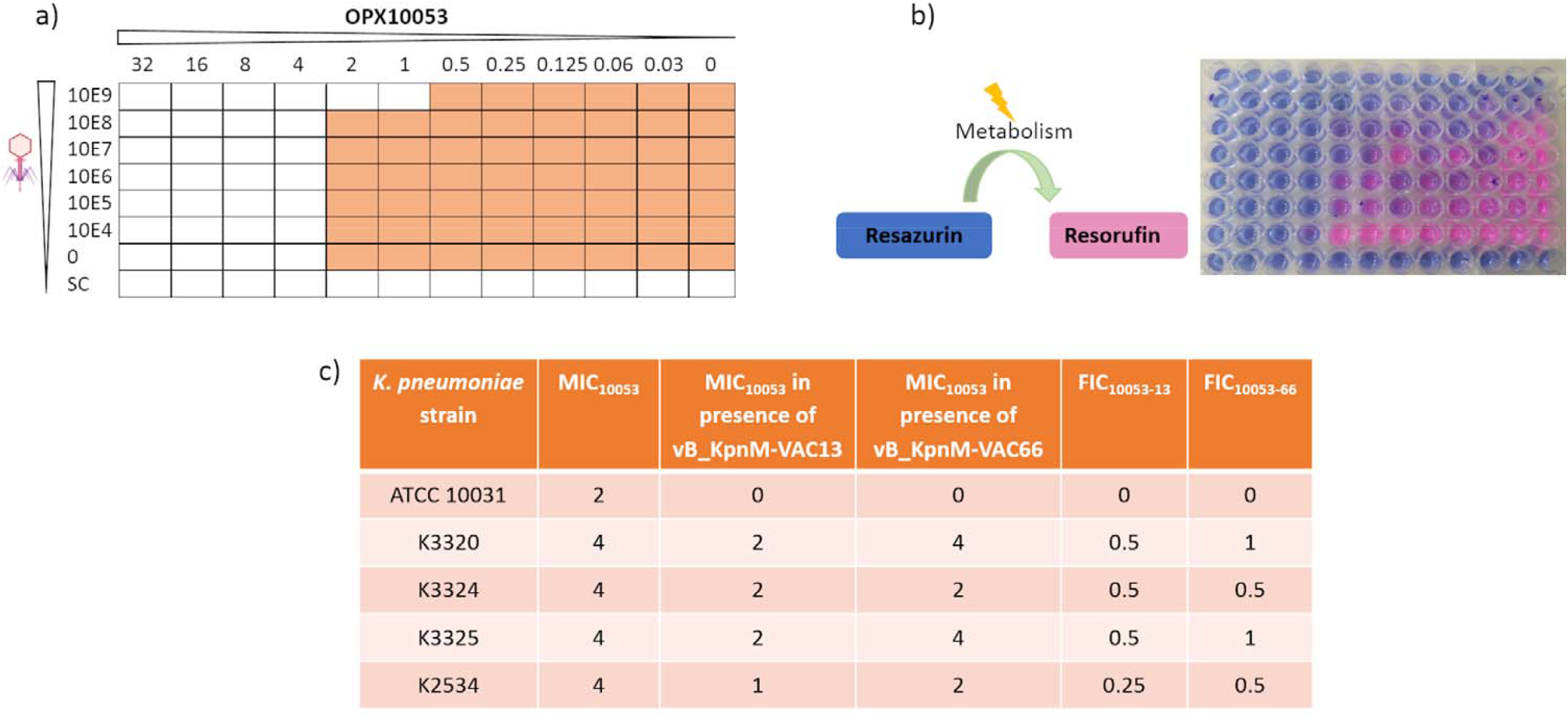
Checkerboard assay in the isolate K2534 (a) and resazurin staining (b) for assessment of metabolically active cells in presence of phages and the octapeptin OPX10053. A representative plate is shown. c) Fractional inhibitory concentration (FIC) of OPX10053 in presence of lytic phages vB_KpnM-VAC13 and vB_KpnM-VAC66. All MIC values are expressed in μg/mL

### Optical density growth curves

Considering the checkerboard assays and the FIC calculations, we used clinical isolates K3324 and K2534 together with the reference strain ATCC 10031 to confirm the synergistic effect between the lytic phages and the peptide OPX10053. According to the infection kinetics (Figure 2) and the previous determination of a considerably high frequency of resistance mutants for both phages ^26, 34^, we decided to sequentially apply the phage and lipopeptide agents. We first exposed the bacterial cells to each bacteriophage for 4h, as we determined this time-point to be the earliest in the appearance of resistance, followed by the addition of OPX10053 at a ½ MIC, providing a final concentration of 2 μg/mL (for K3324 and K2534) or 1 μg/mL (ATCC 10031), with the OD_600 nm_ being measured hourly. The growth of K3324 was similarly inhibited by the phage vB_KpnM-VAC13 alone compared to the combination of this phage with OPX10053; the same phenomenon was not true for the phage vB_KpnM-VAC66, for which a regrowth started from 11 hpi onwards; this was inhibited with the combined effect of vB_KpnM-VAC66 and OPX10053 (Figure 4a). Similarly, the combination of vB_KpnM-VAC13+OPX10053 inhibited K2534 growth compared to the respective monotherapies (Figure 4a). The combination of vB_KpnM-VAC66+OPX10053 was not effective against K2534 *in vitro*. Finally, testing against reference strain ATCC 10031 revealed that both the phages alone and combined with OPX10053 produced a drastic reduction in the OD_600 nm_ of the population, suggesting a high susceptibility of this strain towards the lytic activity of vB_KpnM-VAC13 and vB_KpnM-VAC66 with no generation of resistance (Figure 4a).

**Figure 4:**
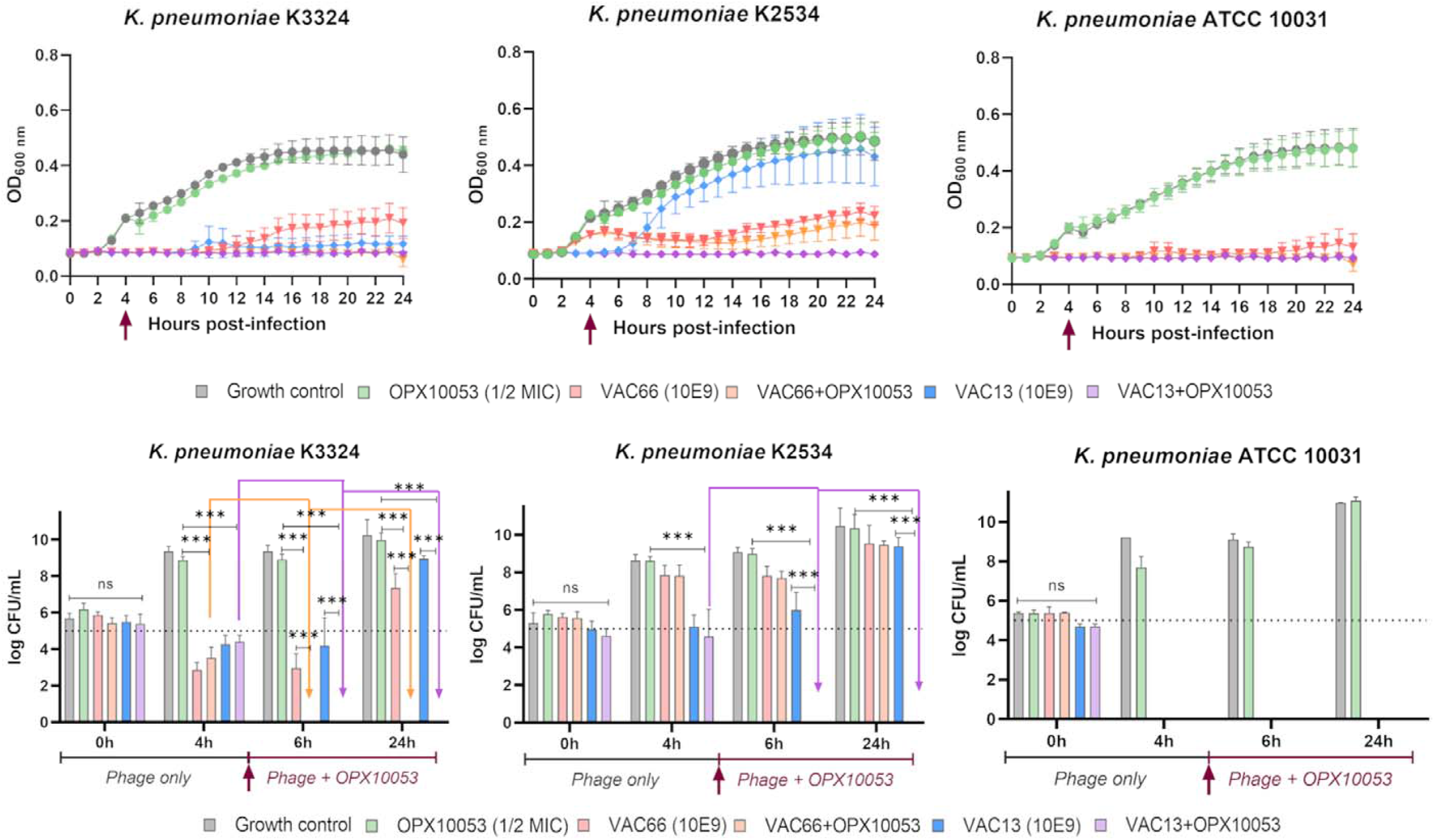
a) Optical density growth curves after combination with both vB_KpnM-VAC13 and vB_KpnM-VAC66 phages and the octapeptin OPX10053. B) Time kill assay after combination with both vB_KpnM-VAC13 and vB_KpnM-VAC66 phages and the octapeptin OPX10053 on K3324, K2534 and ATCC 10031. The dark red arrow indicates the addition of the compound OPX10053 to the medium. ***: p-value <0.001. All the assays were performed in triplicate and statistically analysed with GraphPrism 9.0.

### Time kill assay

A time-kill assay was performed in order to assess the reduction in the viability of bacterial cultures *in vitro*. The bacterial counts (colony forming units per mL, or CFU/mL) were enumerated at 0, 4, 6 and 24 hpi, and the octapeptin OPX10053 was added after the 4h-enumeration. Regarding the effect of vB_KpnM-VAC13, we observed a 5-log/4-log reduction in the viability of K3324 and K2534 strains, respectively, after 4 hpi when compared to the control or the OPX10053 alone (Figure 4b, blue bars); focusing on vB_KpnM-VAC66, a 6-log reduction in the K3324 CFUs was assessed, in contrast to the absence of a statistically significant decrease in the case of K2534 (Figure 4, light red bars). Most importantly, combinations of both phages with OPX10053 reduced the CFU counts of both isolates to as low as nearly 0 for K3324 (represented by orange and purple arrows in Figure 4b). Moreover, vB_KpnM-VAC13+OPX10053 produced this same effect on K2534 strain (purple arrow in Figure 4b), and this synergy was maintained until 24 hpi, as depicted in Figure 4b. The fact that this decrease lasted until 24 hpi suggests that no resistant mutants could regrow. No synergistic activity for any bacteriophage in combination with the compound OPX10053 was observed for the reference isolate ATCC 10031, highlighting the dominant effect of the phages alone.

### Toxicity and efficacy assay: *G. mellonella* infection model

To determine if the octapeptin OPX10053 alone and combined with lytic phages at subinhibitory concentrations was toxic and effective *in vivo, G. mellonella* larvae were injected via their last left proleg with this compound and their survival was monitored for 48h (Figure 5). Interestingly, when injected alone or combined with phages, OPX10053 was found to be toxic, exhibiting high rates of mortality (Figure 5). In contrast, non-infected larvae injected with a solution containing exclusively phages exhibited the same mortality rates as the ones injected with saline, proving absence of toxicity. As shown in Figure 5 b, c, d and e, the phages alone protected the larvae infected with either K3324 or K2534 in a statistically significant way compared to the infection control group (p-values 0.019 and 0.0001 with vB_KpnM-VAC13 and vB_KpnM-VAC66 for K3324, and p-values 0.005 and 0.008 for K2534). For K3324 isolate, vB_KpnM-VAC13 alone protected against the combined treatment (p-value 0.027), just as vB_KpnM-VAC66 did (p-value 0.0006). This latter also conferred protection compared to the OPX10053 alone (p-value 0.0006).

**Figure 5:**
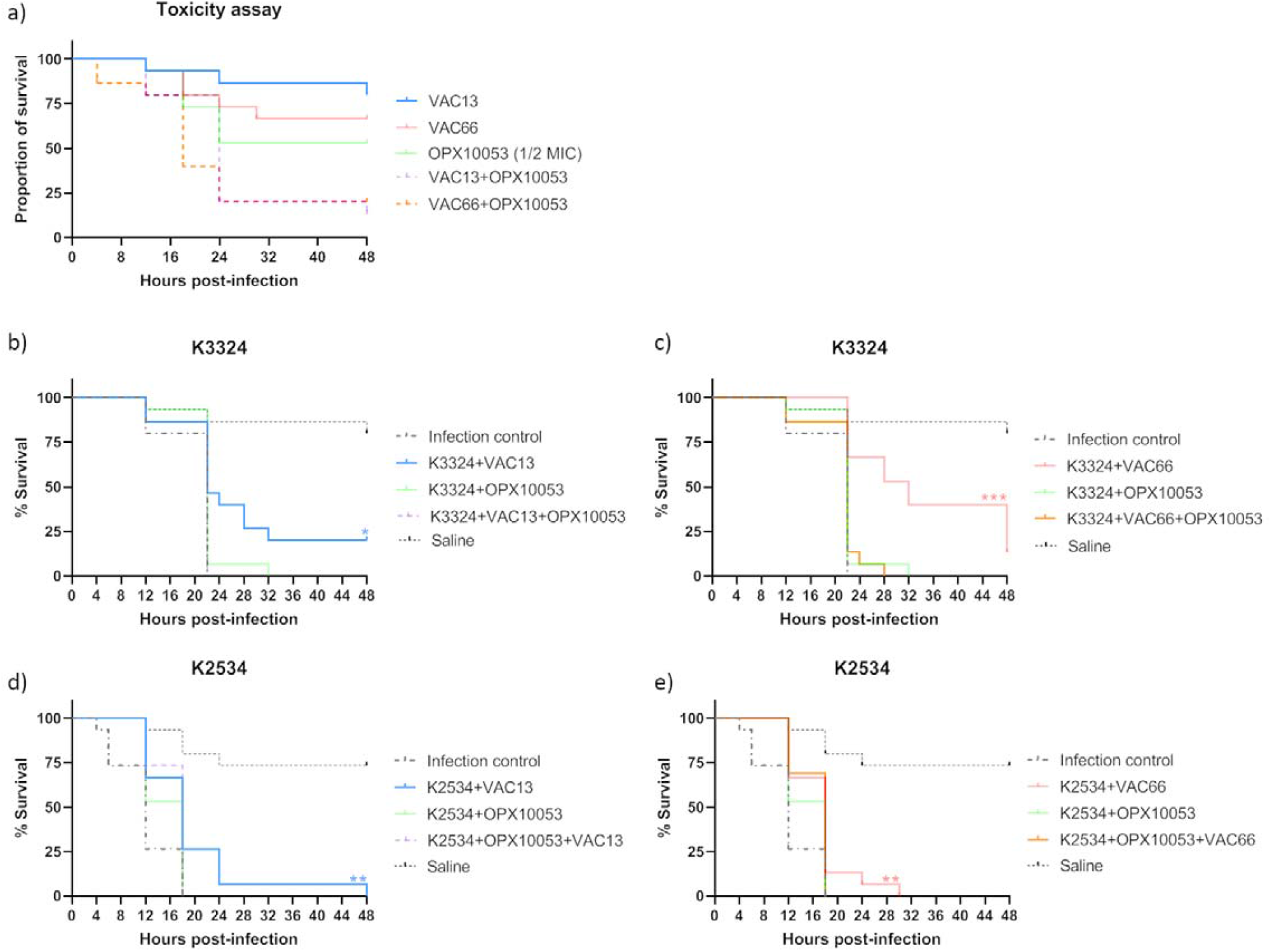
*Galleria mellonella* toxicity and efficacy assays. Kaplan-Meier curves indicate the percentage of survival; the statistical significance was assessed using the Mantel-Cox test (GraphPrism 9.0), where * indicates p-value <0.05, ** corresponds to p-value <0.01 and *** p-value <0.001.

## Discussion

In the actual crisis of increasing antimicrobial resistance, combinations of two or more antimicrobial agents could become a good strategy to counteract infections ^23^. The renewed interest in lytic phages as a possible therapeutic solution has one major drawback, which is the highly resistant profiles that bacteria display to them.

By combining lytic phages with the octapeptin OPX10053, we observed a synergistic effect assessed by MIC and FIC calculations, growth curves and time-kill assay. For K3324, both vB_KpnM-VAC13 and vB_KpnM-VAC66 synergized with the octapeptin OPX10053 *in vitro*, whereas for K2534 this synergy was only found for the former bacteriophage, probably due to its ability to establish a very fast infection of this strain, as previously characterized ^34^. We hypothesized that the reason for this synergy is that preliminary exposure to phages drastically reduces bacterial density, with the sequential addition of octapeptin preventing the rise of resistant mutants. Similar works combining lytic phages (PEV20, an anti-*P. aeruginosa* phage) with amikacin, colistin and ciprofloxacin also showed synergism against highly phage-susceptible isolates whereas no synergistic effect was achieved in the lowest phage-susceptible strain. Akturk *et al*. found that the simultaneous addition of gentamicin and ciprofloxacin with phages resulted in no synergistic effect against a dual biofilm of *P. aeruginosa* and *S. aureus*, whereas sequential addition led to significant reduction of the biofilm biomass when the antibiotics were added after 6h of phage exposure ^36^. Other works have combined a phage-derived product, an endolysin from an *Acinetobacter baumannii* phage, with colistin (structurally and mechanistically similar to the OPX10053 reported here), with a good therapeutic potential both *in vitro* and *in vivo* ^37^. In a different study, Han *et al*. combined a lytic phage with polymyxin B against *K. pneumoniae* and found a synergistic killing *in vitro* with no cross-resistance events ^28^.

We wanted to assess the efficacy of OPX10053 combined with lytic phages *in vivo*, so we inoculated both agents into *G. mellonella* larvae and monitored their survival. Many articles report the use of this model to validate the efficacy of phages and peptides in the *in vivo* system, as it allows flexibility to test different strains and therapeutic combinations more easily and cheaply than the murine model ^38-42^. Furthermore, the immune systems of invertebrates and humans share some elements concerning the primary innate immune response. In our experiments, the octapeptin OPX10053 did not protect larvae against the two *K. pneumoniae* strains tested. We hypothesized that the peptide was exerting a toxic effect on the larval hemocytes, which could explain the high rates of mortality observed in the groups treated either with OPX10053 alone or in combination with the phages, comparable to the infection control group (Figure 5). Nonetheless, when the larvae were treated with vB_KpnM-VAC13 or vB_KpnM-VAC66 alone, a slight but statistically significant protection was conferred compared to the infection control group and the groups treated with OPX10053 alone and combined with phages (Figure 5).

The most striking protection was conferred by vB_KpnM-VAC66 in the K3324-infected larvae, consistent with the 6-log reduction in the CFU enumerated at 4 hpi with vB_KpnM-VAC66, assessed by the time-kill assay (Figure 4b, light red bars). In contrast, the combination of vB_KpnM-VAC66 and OPX10053 was ineffective in reducing the viability of K2534 *in vivo*, contrary to the drastic synergistic effect between vB_KpnM-VAC13 and OPX10053 in this isolate (Figure 4b). However, a major limitation in the comparison of these two assays is the fact that in the *in vivo* experiment, phages and OPX10053 had to be simultaneously applied, as three injections were not feasible.

To our knowledge, this is the first time that synergism between lytic phages and chemically synthesized octapeptins has been demonstrated in vitro and in vivo. This suggests that phages might increase the susceptibility to octapeptins, which could be used in sub-inhibitory concentrations to promote safer dosing. However, our studies did not show a decrease of the toxicity of octapeptin in combination with phages in G. mellonella model. Even so, this innovative combination approach could open the door to some effective drug-biological entity associations.

## Acknowledgements

We would like to acknowledge Raghu Bolisetti for the synthesis and purification of octapeptin analogues, to Dr Alysha Elliott, Angela Kavanagh and Maite Amado for their assistance with the experiments, and to the Institute for Molecular Biosciences and the University of Queensland for providing a safe space where most of the experiments here related could have been conducted.

## Funding

This study was funded by the grants PI19/00878 and PI22/00323 awarded to M. Tomás within the State Plan for R+D+I 2013-2016 (National Plan for Scientific Research, Technological Development and Innovation 2008-2011) and co-financed by the ISCIII-Deputy General Directorate for Evaluation and Promotion of Research-European Regional Development Fund “A way of Making Europe” and Instituto de Salud Carlos III FEDER, (CIBER, PMP22/00092 and the grant PMP22/00092, Personalized Precision Medicine Project MePRAM), the Study Group on Mechanisms of Action and Resistance to Antimicrobials, GEMARA (SEIMC, http://www.seimc.org/) and PIRASOA program for prevention and control of infections related to health care and appropriate use of antimicrobials in Andalusia. M. Tomás was financially supported by the Miguel Servet Research Programme (SERGAS and ISCIII). I. Bleriot was financially supported by the pFIS program (ISCIII, FI20/00302). O. Pacios, L. Fernández-García and M. López were financially supported by the grants IN606A-2020/035, IN606B-2021/013 and IN606C-2022/002, respectively (GAIN, Xunta de Galicia).

## Transparency declaration

Nothing to declare

## Notes

### Competing Interest Statement

The authors have declared no competing interest.

## References

1. Hitt SJ, Bishop BM, van Hoek ML. Komodo-dragon cathelicidin-inspired peptides are antibacterial against carbapenem-resistant Klebsiella pneumoniae. J Med Microbiol 2020; 69: 1262–72.

2. Bachiri T, Bakour S, Lalaoui R et al. Occurrence of Carbapenemase-Producing Enterobacteriaceae Isolates in the Wildlife: First Report of OXA-48 in Wild Boars in Algeria. Microb Drug Resist 2018; 24: 337–45.

3. Golkar Z, Bagasra O, Pace DG. Bacteriophage therapy: a potential solution for the antibiotic resistance crisis. J Infect Dev Ctries 2014; 8: 129–36.

4. Prevention CfDCa. Antibiotic resistant threats in the United States. 2019.

5. Trastoy R, Manso T, Fernandez-Garcia L et al. Mechanisms of Bacterial Tolerance and Persistence in the Gastrointestinal and Respiratory Environments. Clin Microbiol Rev 2018; 31.

6. Clegg S, Murphy CN. Epidemiology and Virulence of Klebsiella pneumoniae. Microbiol Spectr 2016; 4.

7. Pacios O, Fernández-García L, Bleriot I et al. Adaptation of clinical isolates of Klebsiella pneumoniae to the combination of niclosamide with the efflux pump inhibitor phenyl-arginine-β-naphthylamide (PaβN): co-resistance to antimicrobials. J Antimicrob Chemother 2022; 77: 1272–81.

8. Spapen H, Jacobs R, Van Gorp V et al. Renal and neurological side effects of colistin in critically ill patients. Ann Intensive Care 2011; 1: 14.

9. Justo JA, Bosso JA. Adverse reactions associated with systemic polymyxin therapy. Pharmacotherapy 2015; 35: 28–33.

10. Kelesidis T, Falagas ME. The safety of polymyxin antibiotics. Expert Opin Drug Saf 2015; 14: 1687–701.

11. Roberts KD, Zhu Y, Azad MAK et al. A synthetic lipopeptide targeting top-priority multidrug-resistant Gram-negative pathogens. Nat Commun 2022; 13: 1625.

12. Brown P, Abbott E, Abdulle O et al. Design of Next Generation Polymyxins with Lower Toxicity: The Discovery of SPR206. ACS Infect Dis 2019; 5: 1645–56.

13. Blaskovich MAT, Pitt ME, Elliott AG et al. Can octapeptin antibiotics combat extensively drug-resistant (XDR) bacteria? Expert Rev Anti Infect Ther 2018; 16: 485–99.

14. Becker B, Butler MS, Hansford KA et al. Synthesis of octapeptin C4 and biological profiling against NDM-1 and polymyxin-resistant bacteria. Bioorg Med Chem Lett 2017; 27: 2407–9.

15. Han ML, Shen HH, Hansford KA et al. Investigating the Interaction of Octapeptin A3 with Model Bacterial Membranes. ACS Infect Dis 2017; 3: 606–19.

16. Velkov T, Gallardo-Godoy A, Swarbrick JD et al. Structure, Function, and Biosynthetic Origin of Octapeptin Antibiotics Active against Extensively Drug-Resistant Gram-Negative Bacteria. Cell Chem Biol 2018; 25: 380-91.e5.

17. Pitt ME, Cao MD, Butler MS et al. Octapeptin C4 and polymyxin resistance occur via distinct pathways in an epidemic XDR Klebsiella pneumoniae ST258 isolate. J Antimicrob Chemother 2019; 74: 582–93.

18. Zabawa TP, Pucci MJ, Parr TR et al. Treatment of Gram-negative bacterial infections by potentiation of antibiotics. Curr Opin Microbiol 2016; 33: 7–12.

19. Qian CD, Wu XC, Teng Y et al. Battacin (Octapeptin B5), a new cyclic lipopeptide antibiotic from Paenibacillus tianmuensis active against multidrug-resistant Gram-negative bacteria. Antimicrob Agents Chemother 2012; 56: 1458–65.

20. Gordillo Altamirano FL, Barr JJ. Phage Therapy in the Postantibiotic Era. Clin Microbiol Rev 2019; 32.

21. Chan BK, Abedon ST, Loc-Carrillo C. Phage cocktails and the future of phage therapy. Future Microbiol 2013; 8: 769–83.

22. Dabrowska K, Abedon ST. Pharmacologically Aware Phage Therapy: Pharmacodynamic and Pharmacokinetic Obstacles to Phage Antibacterial Action in Animal and Human Bodies. Microbiol Mol Biol Rev 2019; 83.

23. Pacios O, Blasco L, Bleriot I et al. Strategies to Combat Multidrug-Resistant and Persistent Infectious Diseases. Antibiotics (Basel) 2020; 9.

24. Mu A, McDonald D, Jarmusch AK et al. Assessment of the microbiome during bacteriophage therapy in combination with systemic antibiotics to treat a case of staphylococcal device infection. Microbiome 2021; 9: 92.

25. Blasco L, Ambroa A, Lopez M et al. Combined Use of the Ab105-2phiDeltaCI Lytic Mutant Phage and Different Antibiotics in Clinical Isolates of Multi-Resistant Acinetobacter baumannii. Microorganisms 2019; 7.

26. Pacios O, Fernández-García L, Bleriot I et al. Phenotypic and Genomic Comparison of Klebsiella pneumoniae Lytic Phages: vB_KpnM-VAC66 and vB_KpnM-VAC13. Viruses 2021; 14.

27. Maffei E, Shaidullina A, Burkolter M et al. Systematic exploration of Escherichia coli phage-host interactions with the BASEL phage collection. PLoS Biol 2021; 19: e3001424.

28. Han ML, Nang SC, Lin YW et al. Comparative metabolomics revealed key pathways associated with the synergistic killing of multidrug-resistant Klebsiella pneumoniae by a bacteriophage-polymyxin combination. Comput Struct Biotechnol J 2022; 20: 485–95.

29. Jiang X, Patil NA, Azad MAK et al. A novel chemical biology and computational approach to expedite the discovery of new-generation polymyxins against life-threatening Acinetobacter baumannii. Chem Sci 2021; 12: 12211–20.

30. Leclercq R, Cantón R, Brown DF et al. EUCAST expert rules in antimicrobial susceptibility testing. Clin Microbiol Infect 2013; 19: 141–60.

31. Engeman E, Freyberger HR, Corey BW et al. Synergistic Killing and Re-Sensitization of Pseudomonas aeruginosa to Antibiotics by Phage-Antibiotic Combination Treatment. Pharmaceuticals (Basel) 2021; 14.

32. Aradhana V, Srividya ND, Raghu PJ et al. Determining the Minimum Inhibitory Concentration of Bacteriophages: Potential Advantages. Advances in Microbiology, 2013.

33. Bonilla N, Rojas MI, Netto Flores Cruz G et al. Phage on tap-a quick and efficient protocol for the preparation of bacteriophage laboratory stocks. PeerJ 2016; 4: e2261.

34. Pacios O, Fernández-García L, Bleriot I et al. Enhanced Antibacterial Activity of Repurposed Mitomycin C and Imipenem in Combination with the Lytic Phage vB_KpnM-VAC13 against Clinical Isolates of Klebsiella pneumoniae. Antimicrob Agents Chemother 2021; 65: e0090021.

35. Odds FC. Synergy, antagonism, and what the chequerboard puts between them. J Antimicrob Chemother 2003; 52: 1.

36. Akturk E, Oliveira H, Santos SB et al. Synergistic Action of Phage and Antibiotics: Parameters to Enhance the Killing Efficacy Against Mono and Dual-Species Biofilms. Antibiotics (Basel) 2019; 8.

37. Blasco L, Ambroa A, Trastoy R et al. In vitro and in vivo efficacy of combinations of colistin and different endolysins against clinical strains of multi-drug resistant pathogens. Sci Rep 2020; 10: 7163.

38. Tkhilaishvili T, Wang L, Tavanti A et al. Antibacterial Efficacy of Two Commercially Available Bacteriophage Formulations, Staphylococcal Bacteriophage and PYO Bacteriophage, Against Methicillin-Resistant Staphylococcus aureus : Prevention and Eradication of Biofilm Formation and Control of a Systemic Infection of Galleria mellonella Larvae. Front Microbiol 2020; 11: 110.

39. Tsai CJ, Loh JM, Proft T. Galleria mellonella infection models for the study of bacterial diseases and for antimicrobial drug testing. Virulence 2016; 7: 214–29.

40. Thiry D, Passet V, Danis-Wlodarczyk K et al. New Bacteriophages against Emerging Lineages ST23 and ST258 of Klebsiella pneumoniae and Efficacy Assessment in Galleria mellonella Larvae. Viruses 2019; 11.

41. Forti F, Roach DR, Cafora M et al. Design of a Broad-Range Bacteriophage Cocktail That Reduces Pseudomonas aeruginosa Biofilms and Treats Acute Infections in Two Animal Models. Antimicrob Agents Chemother 2018; 62.

42. Pompilio A, Geminiani C, Mantini P et al. Peptide dendrimers as “lead compounds” for the treatment of chronic lung infections by Pseudomonas aeruginosa in cystic fibrosis patients: in vitro and in vivo studies. Infect Drug Resist 2018; 11: 1767–82.

